# Liver Sinusoidal Endothelial Cells and Laminin dictate cholangiocytes’ fate in chronic liver disease

**DOI:** 10.1101/2024.09.29.615650

**Authors:** Rita Manco, Camilla Moliterni, Gauthier Neirynck, Maxime De Rudder, Corinne Picalausa, Leana Ducor, Montserrat Fraga, Frédéric Lemaigre, Christine Sempoux, Alexandra Dili, Isabelle A. Leclercq

## Abstract

In chronic liver disease (CLD), hepatocyte senescence limits conventional regeneration, prompting an alternative regeneration mechanism through transdifferentiation of reactive cholangiocytes (or ductular reaction cells - DRs). Yet, this alone is insufficient to avoid liver failure and transplantation, highlighting the importance of a deeper understanding of the mechanism. We focused on the DRs microenvironment. We identified laminin as a crucial component of the extracellular matrix to maintain DRs in a biliary phenotype, preventing their differentiation into hepatocytes. Liver sinusoidal endothelial cells (LSEC) are the cells regulating laminin degradation around DRs. By preventing LSEC capillarization during CLD, we enhanced DR differentiation into hepatocytes. We also demonstrated this causality in human samples. This is the first time that a mechanism for DR-driven regeneration has been described.

## Introduction

The liver possesses a high capacity to regenerate. Due to its versatility, the liver adopts different regenerative strategies depending on the underlying damage^1^. Following partial hepatectomy and acute liver injury, regeneration occurs through a well-orchestrated mechanism with replication of each individual cell type^2^. By contrast in the context of long-lasting liver disease, hepatocytes lose their proliferative capacity entering the so-called replicative senescence^3^. The accumulation of senescent hepatocytes hampers the ‘canonical’ mechanism of liver regeneration, resulting in a decline in hepatic function and decompensation. In response to this scenario, an alternative mechanism of regeneration is initiated^4–6^. This mechanism produces new *bona fide* hepatocytes by differentiation of reactive cholangiocytes – also called ductular reaction cells (DR). The DR-derived hepatocytes, originating from a single cholangiocyte, undergo clonal expansion, more resistant to DNA damage compared to native hepatocytes, and demonstrate reduced susceptibility to carcinogenesis stimuli^6^. Despite these advantages, this alternative regeneration mechanism alone is insufficient for liver function recovery. Consequently, patients suffering from chronic liver disease may eventually require liver transplantation. The question arises as to whether DR-driven regeneration can be enhanced for therapeutic purposes, highlighting the importance of a deeper understanding of the mechanisms.

Cholangiocytes, also called biliary epithelial cells (BECs), are mature epithelial cells that line the bile ducts^7^. BECs perform liver-specific functions, including the modification and transport of the bile. They exhibit considerable plasticity, making them a reservoir of liver progenitor cells that undergo dynamic changes in response to tissue damage. These changes include exit from a quiescent state and proliferating as DRs, to eventually differentiate into hepatocytes. Historically, BECs were classified based on their morphology and functional properties with changes along the biliary tree, from small to large ducts^8^. According to this classification, the differentiation capacity was thought to predominantly reside in the BECs of the small ducts, possessing a more immature phenotype. Other researchers even suggest the existence of a true bipotential progenitor able to differentiate into BECs or hepatocytes according to injury context. However, such a population was not confirmed in more recent studies using single-cell RNA sequencing, which attributes to all BECs the same capacity in response to injury^9,10^. The response to injury was driven by the Hippo-YAP pathway, thus leaving to the microenvironment the lead in orienting the BECs’ fate. In performing the main liver-specific functions, liver epithelial cells are assisted by the so-called non-parenchymal cells (NPCs) that provide a structure as well as establish contact and collaboration with them. Liver sinusoidal endothelial cells (LSECs) account for approximately 50% of NPCs and have contact with both hepatocytes and cholangiocytes. LSECs in healthy liver exhibit unique features and functions, including fenestration, and lie on a discontinuous basement membrane^11^. These particularities allow a bidirectional free exchange between blood and hepatocytes. They dictate the spatial distribution of labor in the hepatocytes along the lobule axis^12^. LSECs also prevent the activation of the hepatic stellate cells (HSCs), hence controlling fibrotic remodeling. The loss of fenestration and the thickening of the basement membrane, a phenomenon named “capillarization”, is considered to be a key event of the pathological remodeling of the lobular architecture during liver disease^13^. Capillarized LSECs promote the activation of HSCs and deposition of ECM, resulting in fibrosis. Intervention to reverse LSECs capillarization supports the deactivation of HSC and regression of fibrosis^14^. The role of LSECs in DR-driven regeneration remains unknown. We therefore investigated the role of the microenvironment during DR-driven regeneration in a mouse model of chronic liver disease. We used animal models, imaging analysis, -omics technologies, in vitro double spheroids to demonstrate that LSEC phenotype shapes the microenvironment around DRs and guide their fate. The operation of such a mechanism is supported by spatial observations in human diseased livers.

### Laminin as gatekeeper of the biliary phenotype

The literature supports that all BECs exhibit the same capacity to respond to injury via the Hippo-YAP pathway^9,10^. The latter has been identified as mechanotransducers^15^. The basal side of BECs in the bile ducts is in close contact with the basement membrane, primarily composed of laminin as the most abundant non-collagenous ECM component^16^ and Collagen IV ^17^. To follow the fate of BEC/DRs in DR-driven regeneration, we used the osteopontin-iCreER^T2^ (OPN-Cre) mice crossed with the Rosa26^YFP^. In this model, upon tamoxifen injection and in the course of the CCl_4_ treatment, YFP expression is restricted to the BECs, DRs, and DR-derived hepatocytes^6,18^. As for note, we refer to DRs as at least one or more CK19^+^ or YFP^+^ cells in line and extending out of the bile duct, into the parenchyma (See Fig. 1A, C – Lam-DR and NoLam-DR). In livers from mice repeatedly injected with CCl_4_ for 6 weeks, we identified two distinct groups of YFP^+^ DRs: those surrounded by a laminin layer (Lam-DRs) and those not surrounded by laminin (noLam-DRs) (Fig. 1A-B). NoLam-DRs represent ∼20% of the entire DRs population (Fig. 1B). Notably, in the CCl_4_ disease model we did not observe differences in DRs populations when stained for Collagen I and Collagen IV, which was not detected around DRs (Fig. S1).

**Fig. 1.**
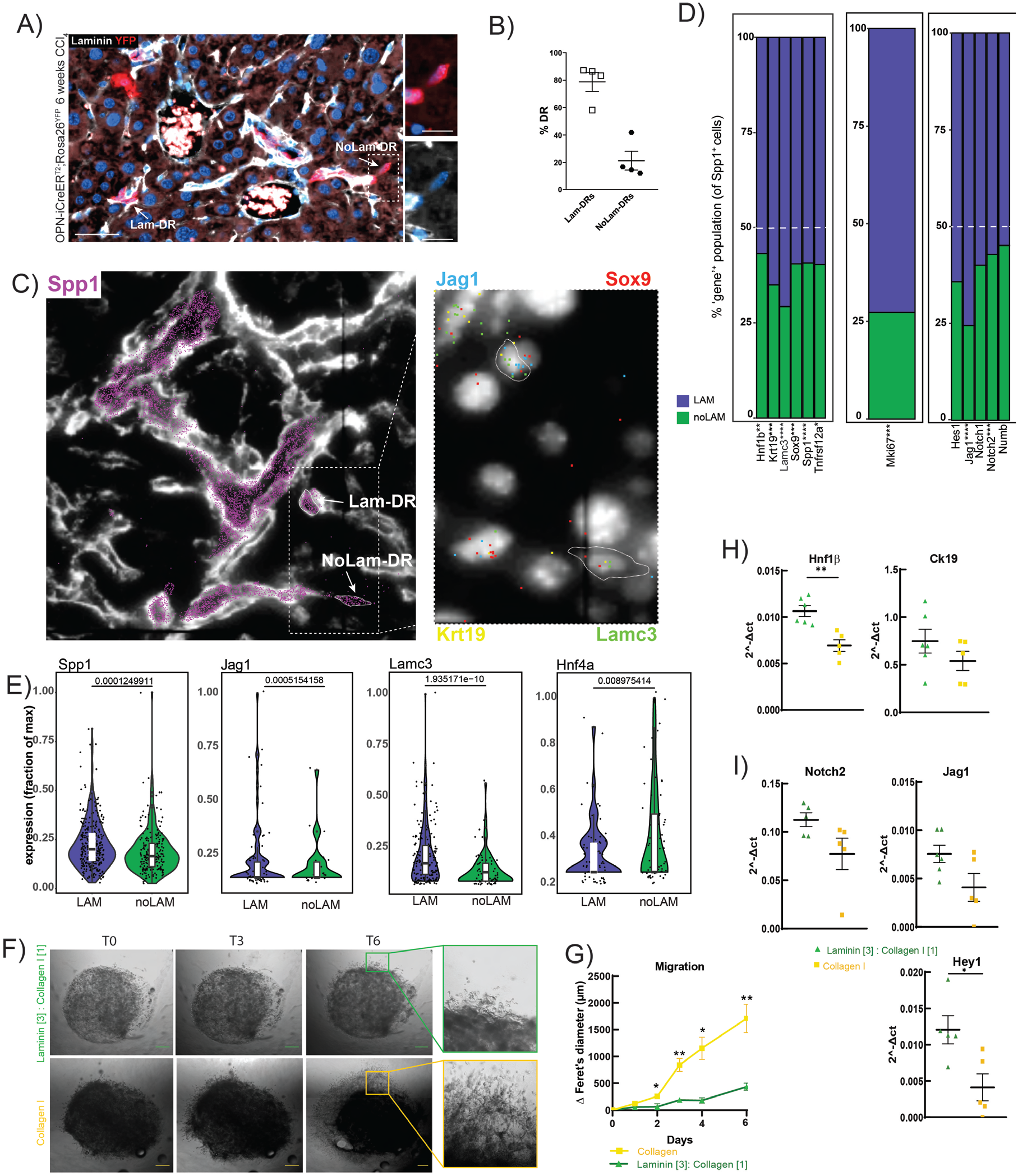
Laminin is the gatekeeper of cholangiocytes phenotype in DR cells. (**A**) Representative immunofluorescent image for Laminin (white) and YFP (red) on liver section from OPN-iCreER^T2^;Rosa26R^YFP^ mice treated for 6 weeks with CCl_4_ (Scale Bar: 50µm). (**B**) Graph showing the quantification of the DRs surrounded or not by laminin in OPN-iCreER^T2^;Rosa26R^YFP^ mice treated for 6 weeks with CCl_4_ (n=4). (**C**) Molecular Cartography of indicated genes and IHC of laminin. (**D**) Bar plots showing the percentage of Spp1+ cells Lam and noLam expressing the indicated gene. p-value was calculated by test of proportion. (**E**) Violin plots of the quantification of the indicated gene in the two populations of DRs, Lam, and noLam. p-value was calculated using the Wilcoxon test. (**F**) Representative photos of the double-spheroids with NMC and Laminin:Collagen I [3:1] (green) or Collagen I (yellow) at different days (T0 – T3 – T6) (scale bar: 200µm ) (**G**) Quantification of the Feret’s diameter as a proxy of the cell migration calculated as Δ on the T0. p-value was calculated using the t-test. (**H**) RT-qPCR of cholangiocyte-specific genes and (**I**) genes of the Notch pathway in the double-spheroids. p-value was calculated using the t-test.

Despite expanding during chronic liver disease, DR cells represent only 0.3% of the total liver area^6^, resulting in a limited number of DRs around each portal triad. Additionally, there are currently no known markers for isolating DRs exclusively. Thus, to interrogate the two distinct DRs groups, Lam-DRs and noLam-DRs, *in vivo*, we employed spatial transcriptomic from Molecular Cartography^TM^ (Resolve BioSciences) enabling 100-plex spatial mRNA analysis. We identified BECs using *Spp1* gene (OPN protein) and segregated them between Lam and NoLam based on the laminin staining (Fig. 1C). Subsequently, we examined both groups for the expression of specific cholangiocytes’ genes (*Hnf1β, Krt19, Sox9, Spp1, Tnfrsf12a, Lamc3*), for genes involved in the Notch pathway (*Hes1, Jag1, Notch1, Notch2, Numb*) as well as the proliferation marker gene *Mki67*. In Figure 1D, we show the percentage of *Spp1*^+^ cells that do express the gene of interest. Compared to Lam-DRs, there is a lower proportion of noLam-DRs that expresses cholangiocytes-specific genes, cell proliferation markers, or genes of the Notch pathway (Fig. 1D). Furthermore, for *Spp1, Jag1*, and *Lamc3* the noLam-DR cells expressing these genes had a lower expression level (Fig. 1E). Cells in the noLam-DRs group demonstrated a significantly higher expression of *Hnf4α* gene (Fig. 1E), the key transcription factor associated with hepatocytes differentiation.

To assess the impact of the laminin layer on the fate of DRs, we cultured Normal Mouse Cholangiocytes (NMC) cells in a double spheroids model ^19^. In this setup, the core comprises a mixture of cholangiocytes and ECM, while the external layer consists of a cell-free ECM. This technology enabled us to precisely examine the influence of ECM proteins on the migration and differentiation of DRs. Spheroids using laminin alone would require the addition of a polymer as a crosslinker that would have affected cell behavior. Therefore, we prepared spheroids with Collagen I only or with a mix of Collagen I and Laminin in a 1:3 ratio. When cultured in a milieu rich in laminin (1:3 collagen: laminin), the cholangiocytes migrate significantly less out of the core than in doubled spheroids composed of collagen I alone, where cells crawl out the core to the ECM ring (Fig. 1F-G). Additionally, in the presence of laminin, cholangiocytes maintain higher expression levels of cholangiocyte-specific genes such as *Ck19* and *Hnf1β* (Fig. 1H) as well as *Notch2, Jag1*, and *Hey1* genes (Fig. 1I) compared to those cultured in collagen I. Conversely, they showed a slight decrease in the expression of hepatocyte-specific genes *Cebpa, Hnf1α*, and *Dbp*, while *Hnf4α* expression remains unchanged (Fig. S2).

The literature reveals a correlation between the presence of the basement membrane and the differentiation of cholangiocytes into hepatocytes ^20,21^. Our data suggest that among the ECM components of the basement membrane, laminin plays a pivotal role in maintaining DRs in a biliary phenotype.

### CD34^+^ LSECs prevent the degradation of laminin around DRs

Senescent cells secrete a wide variety of factors, a phenomenon named senescence-associated secretory phenotype (SASPs). As during a chronic disease the liver becomes senescent, we analyzed the common SASPs factors^3^ and identified a dysregulation of genes associated with angiogenic process, such as *Hgf, Fgf7, Vegfa, and Cxcl12* (Fig. S3A). Also, we stained liver slides of 6 weeks CCl_4_-treated OPN-cre; Rosa^YFP^ for YFP (BECs-DRs), CD31 (endothelial cells) and αSMA (hepatic stellate cells) and found that the cells in closer proximity to DRs were the endothelial cells (Fig. S3B-C). Thus, we decided to investigate further the role of LSECs on DR-driven regeneration.

We focused on the LSECs phenotype. CD34 serves as a marker of capillarization and dysfunctional LSECs^22^. We found that ∼50% of CD34^+^ LSECs were in contact with Lam-DRs, while less than 20% of CD34^+^ LSECs were in contact with noLam-DRs (Fig. 2A-B). Using the spatial transcriptomic data, we demonstrated that LSECs in contact with Lam-DRs exhibit higher expression levels of *Pecam1, Cdh5*, and *Clec4g* (Fig. 2C, Fig. S4), genes upregulated in capillarized LSECs. Also, a higher percentage of LSECs closer to the Lam-DRs express pro-fibrotic genes *Bmp2, Bmp4*, and *Pdgfb* (Fig. 2D). We also analyzed the single-cell RNA sequencing of LSECs sorted from chronically injured mouse livers^23^. When we computationally separated them for *Cd34* expression, we observed that although the two populations of LSECs, *Cd34-* and *Cd34+*, displayed similar expression of MMPs genes (aver. expr. 0.28 vs 0.27), the *Cd34+* population has a higher expression of Laminin genes (aver. expr. 0.48 vs 0.60) as well as Timps genes (aver. expr.4.21 vs 5.14) (Fig. 2E-F, Fig. S5). In an acute injury, CXCR7^+^ LSECs trigger the Id1-mediated production of angiocrine factors, such as Wnt2, contributing to pro-regenerative remodeling^24^. However, during chronic disease, LSECs undergo a shift from CXCR7 to CXCR4 expression, stimulating the proliferation of hepatic stellate cells and adopting a more pro-fibrotic role^24^. Using flow cytometry, we analyzed the expression of these two receptors in the CD34^+^ and CD34^-^ endothelial cells: in the healthy control liver, a higher proportion of CD34^-^ CD31^+^ endothelial cells did not express either receptor (CD34-:53.5% vs CD34+:15.4%, p<0.0001), while CD34^+^CD31^+^ endothelial cells predominantly expressed CXCR7 (CD34-:34.1% vs CD34+:46.5%) (Fig. 2G). After 6 weeks of CCl_4_ treatment, there was a global increase in CXCR4 and in CXCR7 expression, with a higher proportion of these cells being CD34^+^CD31^+^ endothelial cells (for CXCR4: CD34-:46.5% vs CD34+:70.43%, p<0.0001; for CXCR7: CD34-: 72% vs CD34^+^ 95.5%, p<0.0001) (Fig. 2G). This suggests that CD34^+^ LSEC that are preferentially in contact with Lam-DRs adopt a profibrogenic profile compared to the CD34^-^ LSEC in the vicinity of noLam-DRs.

**Fig. 2.**
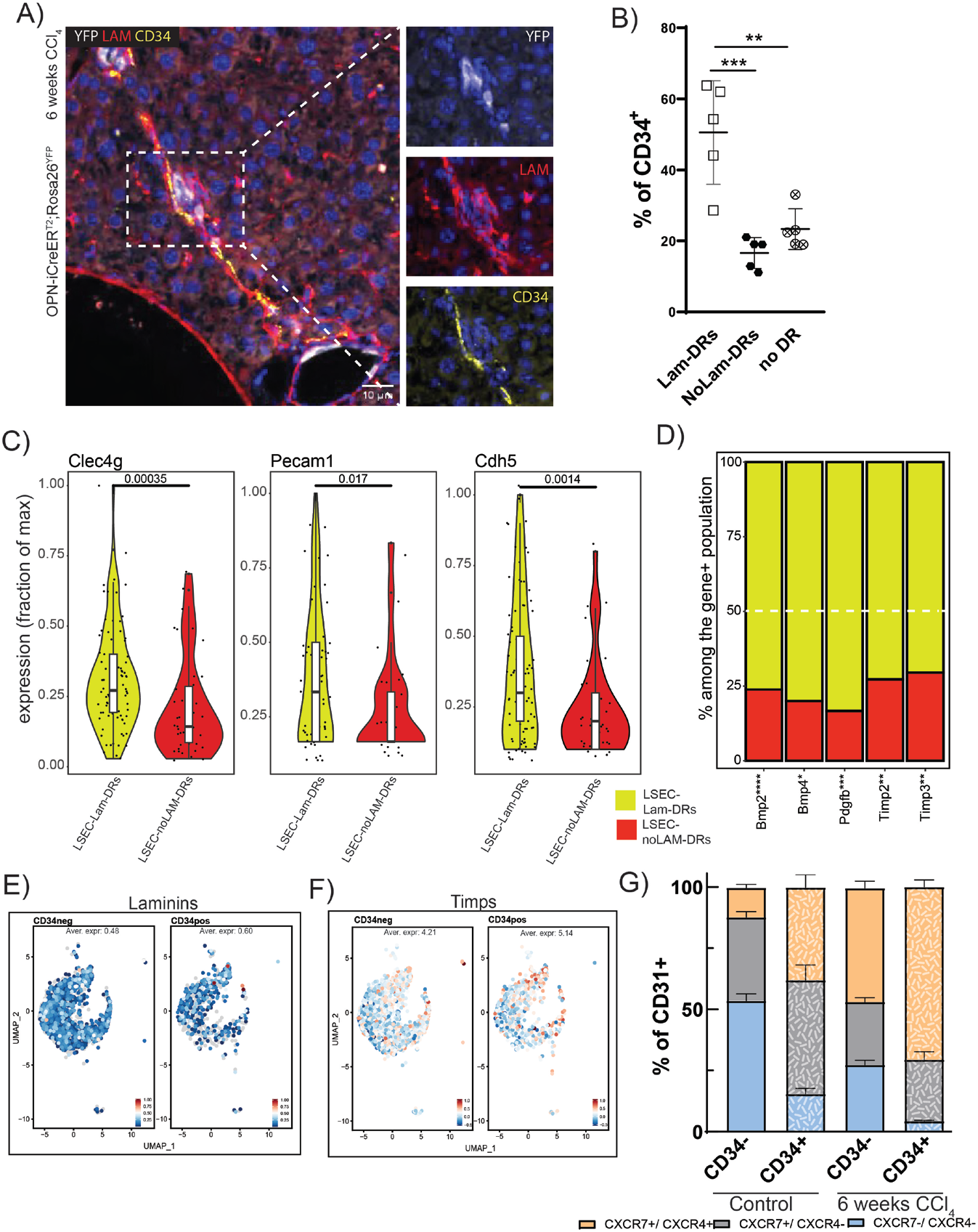
CD34+ LSECs prevent the degradation of laminin around DRs. (**A**) Representative immunofluorescent image for YFP (white), Laminin (red) and CD34 (yellow) on liver section from OPN-iCreER^T2^;Rosa26R^YFP^ mice treated for 6 weeks with CCl_4_ (Scale Bar: 10µm). (**B**) Quantification of CD34+ LSECs closer to Lam-DRs, noLam-DRs or non in contact with DRs. p-value was calculated by 1-way ANOVA with Bonferroni post-test. (**C**) Violin plots of the quantification of the indicated gene in the two populations of LSECs, closer to the Lam-DRs or to the noLam-DRs. p-value was calculated using the Wilcox test. (**D**) Bar plots showing the percentage of LSEC-LamDRs and LSECs-noLamDRs expressing the indicated gene. p-value was calculated by test of proportion. (**E-F**) UMAP plots of LSECs sorted from CCl4-injured mice. Data are from Su et al., cmgh, 2021. (**G**) Stacked barplot of FACS analysis in CD31+ cells of CTL (n=3) and 6weeks CCl4-treated mice (n=6).

Extension of the sinusoidal bed during regeneration after partial hepatectomy is ensured by LSECs proliferation as well as by recruitment and differentiation of bone marrow progenitors^25,26^. To study the origin of LSECs during chronic liver disease we used the Cdh5-Cre; Rosa^mT/mG^ tracing lineage mouse model. In this model, all resident LSECs express the mGreen protein, while any non-resident recruited LSECs are marked as mTomato^+^. Following treatment of the mice with CCl_4_ for 4 and 6 weeks, the percentage of mTomato^+^ recruited cells was similar after CCl_4_ injury than in controls. (Fig. S6).

Overall, our data indicate that during chronic CCl_4_, LSECs emerge as cells in close proximity to DRs. Our results show that CD34^+^CD31^+^ endothelial cells undergo a phenotypic shift from a pro-regenerative state to a more pro-fibrotic profile. This shift impedes the degradation of laminin around DRs, thereby hindering their differentiation into hepatocytes.

### CD34^+^ LSECs – Laminin – DRs complex is also found in human chronic liver diseases

Finally, we investigated if the two groups of DRs, Lam-DRs, and noLam-DRs, were also present in human chronic liver diseases.

Because MASLD/MASH is the most common cause of chronic liver disease, we analyzed the presence or absence of the laminin around DRs in liver biopsies from MASLD/MASH non-cirrhotic patients (Table S1) (Fig. 3A-B). We double-stained the slides for CK7, to visualize DR cells and the DR-derived hepatocytes^27^, and Laminin. For this analysis, we focused on CK7^+^ DRs outside the fibrotic septa. We were able to identify also in human samples the two DRs populations, surrounded (Fig. 3A – blue arrows) or not (Fig. 3A – green arrows) by Laminin.

**Fig. 3.**
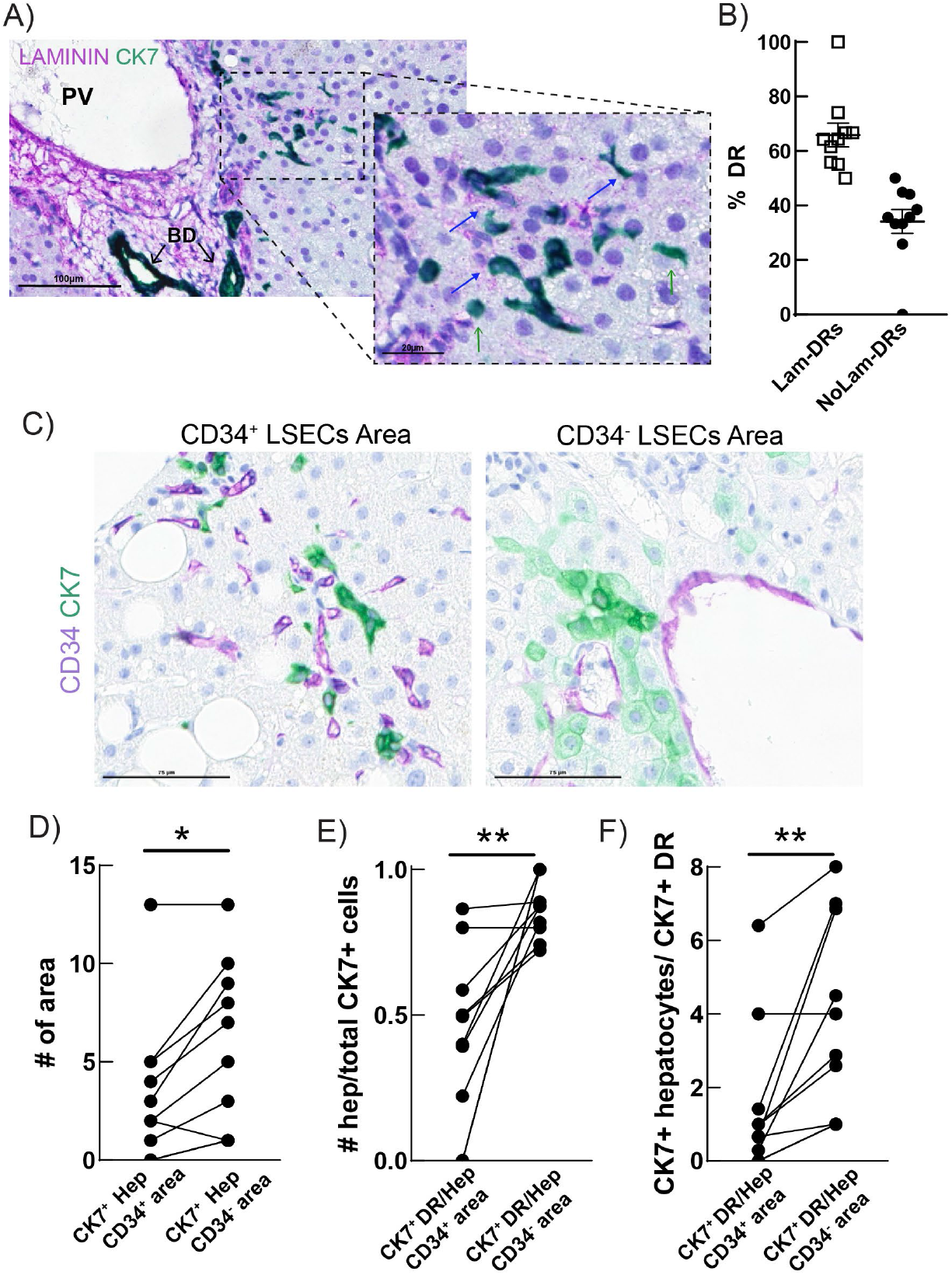
CD34+ LSECs – Laminin – DRs complex in human chronic liver disease. (**A**) Representative images of slides from MASLD patient stained for Laminin (purple) and CK7 (green) showing lam-DRs (blue arrow) and noLam-DRs (green arrow). (**B**) Graph showing the quantification of the DRs surrounded or not by laminin in slides from MASLD patients (n=10) (**C**) Representative images of slides from MASLD patient stained for CD34 (purple) and CK7 (green) showing an area with CD34+ LSECs and CK7+ DRs (left panel) and an area with CD34-LSECs and CK7+ hepatocytes. (**D**) Dot plot representing the number of areas found with CK7+ hepatocytes in CD34+ and CD34-areas. (**E**) Dot plot representing the number of CK7+ hepatocytes over the total CK7+ cells found in CD34+ and CD34-areas. (**F**) Dot plot representing the ratio CK7+ hepatocytes over CK7+ DRs in CD34+ and CD34-areas. P-value was calculated by paired Wilcoxon-test.

Then, we double-stained the samples for CD34, to mark the capillarized LSECs, and again CK7 ^27^. We found CK7^+^ hepatocytes mostly in areas with CD34^-^ LSECs (Fig. 3C, right-hand-sided panel and Fig. 3D). Indeed, within CD34^-^ areas the number of CK7^+^ hepatocytes on the total CK7^+^ cells was higher compared to areas with CD34^+^ LSECs in which CK7+ DR cells were predominant (Fig. 3C, left-hand side panel and Fig. 3E-F). These results suggest that also in human chronic liver disease, CD34^+^ LSECs play a role in maintaining the phenotype of DR cells and impeding their differentiation into new hepatocytes.

We analyzed single-cell LSECs data obtained from MASH without cirrhosis (published by Gribben, Galanakis, et al)^28^. Although the dataset includes data from various stages of the MASL disease spectrum, we focused on MASH without cirrhosis to avoid confounding factors from capillarized endothelial cells located in the septa around the cirrhotic nodules. In the LSECs population there is an overall increase in the number and in the proportion of endothelial cells expressing CD34 (Fig. S7A-B). The CD34^+^ LSECs exhibit higher expression of the Laminin genes (Fig. S7C, Data S2), particularly LAMA1 and LAMA5, which are important components of the cholangiocytes’ basal membrane^16^.

Primary Sclerosis Cholangitis (PSC) is an immune-related cholestatic liver disease. Andrews, Nakib, et al^29^, demonstrated that PSC samples are characterized also by a decrease in HNF4α^+^ cells, an increase in CK7^+^ cells, and importantly an increased number of periportal cells with the co-expression of those two markers^29^. We looked at their single-cell LSECs data and found that only periportal LSECs have an increased expression level of CD34 in PSC (Fig. S8A-B). Similar to the mouse data, CD34^+^ LSECs express higher ACKR3, BMP4, and TIMP4 (Fig. S7C, Data S3), but lower CLEC4G, as also in PSC CD34^+^ LSECs increase the expression of genes to impede laminin remodeling around the DRs. Of note, TIMP4 inhibits MMP-2, MMP-26, and MT1-MMP^30^, all metalloproteinases with higher affinity for the degradation of Laminin^31,32^.

### Preventing LSECs capillarization impacts DR-driven regeneration

As our data pointed to a major role of the capillarized CD34^+^ LSECs in preventing the transdifferentiation of BEC into hepatocytes, we decided to evaluate the importance of the switch between capillarized CD34^+^ and fenestrated CD34^-^ LSECs *in vivo*. To modulate the LSEC capillarization, we used the soluble guanylate cyclase (sGC) activator BAY 60-2770^33^, which has been demonstrated to increase the cGMP activity irrespective of NO binding ^34^. We administrated the compound during the 6-weeks of CCl_4_ treatment in OPN-Cre; RosaYFP mice in two different regimens (Fig. 4A): 1) during the last two weeks of the CCl_4_ treatment to reverse capillarization; and 2) throughout all the 6 weeks of the CCl_4_ treatment to prevent the LSEC capillarization. We were not able to significantly reverse LSECs capillarization in the first experimental setting, but we did prevent it, as shown in the second experimental setting by a significant decrease of the CD34^+^ endothelial cells (Fig. 4B-C). Additionally, we observed less laminin (Fig. 4D) and fewer CK19^+^ cells in the 6w BAY 60-2770 treated group (Fig. 4E). Finally, the YFP^+^ area, signing DR-derived hepatocytes, was significantly increased when applying BAY 60-2770 prevention vs the control group (Fig. 4F-G).

**Fig. 4.**
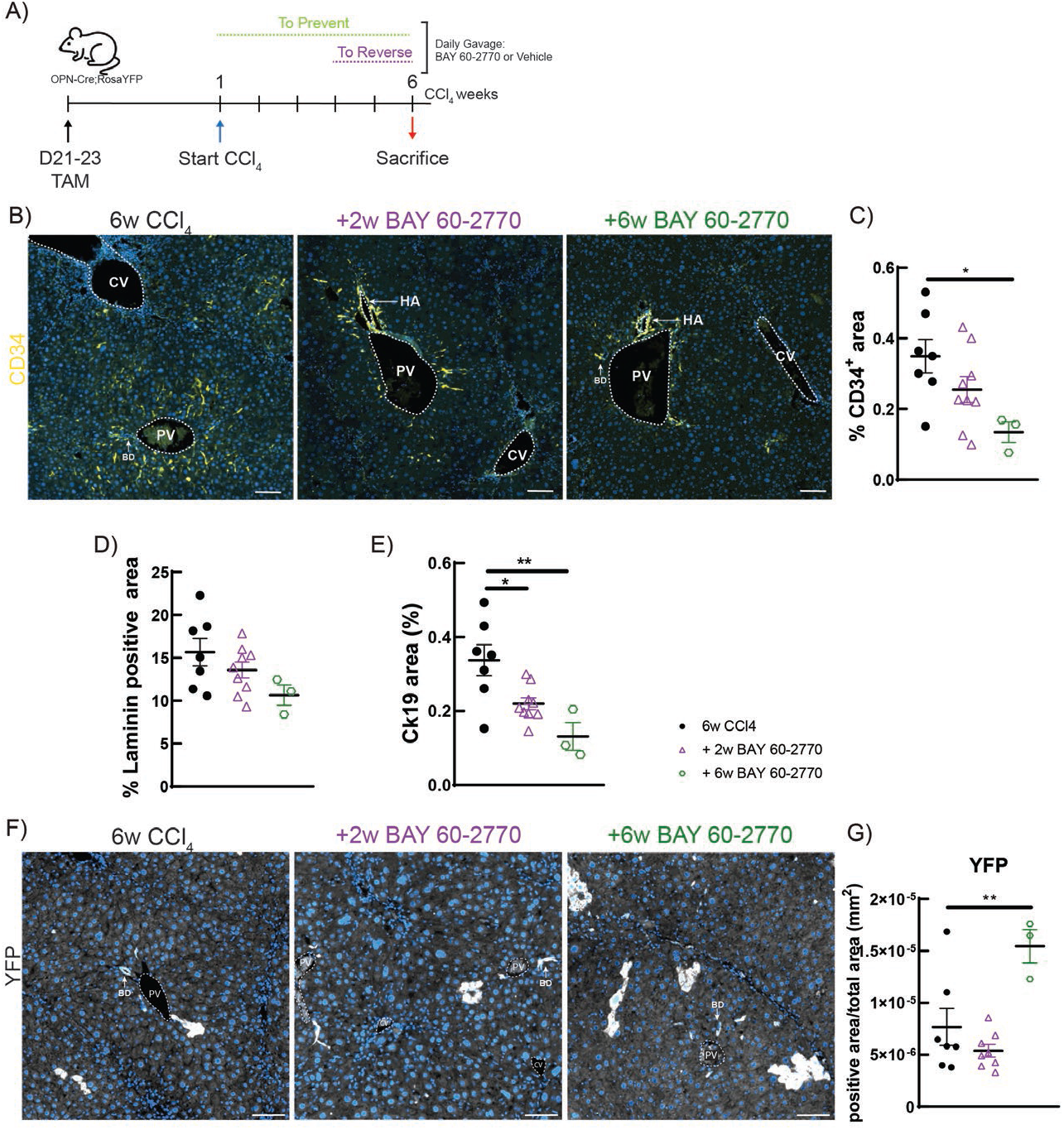
Reverse LSECs capillarization enhances DR-driven regeneration. (**A**)Schematic representation of the experimental design. (**B**) Representative immunofluorescent image for CD34 (yellow) on liver sections from OPN-iCreER^T2^;Rosa26R^YFP^ mice. (**C**) Quantification of CD34+ area in mice treated for 6weeks with CCl4 (n=7), in the ‘reverse’ configuration (n=9) and in the ‘prevent’ configuration (n=3). P-value was calculated by 1-way ANOVA with Bonferroni post-test. (**D**) Quantification of the Laminin positive area in the 3 groups of mice. (**E)** Morphometrical quantification of CK19+ cells. P-value was calculated by 1-way ANOVA with Bonferroni post-test. (**F**) Representative immunofluorescent image for YFP (white) on liver sections from OPN-iCreER^T2^;Rosa26R^YFP^ mice. **(G)** Quantification of YFP+ area in mice treated for 6weeks with CCl4(n=7), in the ‘reverse’ configuration (n=9) and in the ‘prevent’ configuration (n=3). P-value was calculated by 1-way ANOVA with Bonferroni post-test.

## Discussion

DR-driven regeneration emerges as the alternative mechanism of liver regeneration, activated when hepatocytes stop proliferating. Despite holding therapeutic potential, the lack of understanding regarding the mechanism behind DR-driven regeneration limits the development of targeted interventions.

In this study, we proposed for the first time a mechanism that controls the DR-driven regeneration, underscoring the pivotal role played by the LSECs. All BECs have the same capacity to respond to injury, and thus also have the same possibility to differentiate into hepatocytes. This has been convincingly demonstrated by Pepe-Moore et al^9^, and Planas-Paz et al^10^ using single-cell sequencing. We concentrated on investigating the microenvironment around the DRs, as the most likely origin of cues to differentially direct DRs differentiation and fate. We identified two distinct groups of DRs: those that are surrounded by a laminin layer and those that are free of laminin. Laminin potentially blocks DR cells in their ‘biliary’ phenotype as it does for the cholangiocytes of the bile ducts. This is supported by our finding of lower migration of cholangiocytes and higher expression of biliary genes in the presence of laminin. However, in the double spheroids model, the cholangiocytes only slightly increase hepatocytes specific genes when not in contact with laminin. Probably when losing the connection with laminin, DRs become responsive to external signals that prompt their differentiation into hepatocytes, such as Wnt3a secreted by macrophages^35^. Besides ECM, the key difference in the microenvironment of these two DRs populations was the proximity to different LSEC phenotypes: capillarized CD34+ LSECs were closer to Lam-DRs, while fenestrated CD34-LSECs were near the NoLam-DRs. Capillarized CD34+ LSECs expressed higher level of Timps and Lams genes, as well as pro-fibrotic genes. The CD34+ LSECs – Laminin – DRs complex was also found in human chronic liver samples. Finally, preventing the capillarization of the LSECs impacted DR-driven regeneration, resulting in more DR-derived hepatocytes into the liver parenchyma of chronically injured mice.

Senescent hepatocytes do play a role in DR-driven regeneration. Extensive studies demonstrated that replicative senescence in a majority of hepatocytes is a *conditio sine qua non* for DR-sustained parenchyma regeneration^4–6^. When cells become senescent, they acquire a new secretory phenotype (SASP) and start to produce a variety of proteins, including chemokine, interleukin, growth factor, proteases and ECM proteins. Analyzing an extensive list of SASPs secreted molecules we found a dysregulation of angiocrine factors, which prompted us to focus our study on the LSECs. Moreover, when we investigated the DRs immediate microenvironment, we found that LSECs were the most prominent cell population. However, it remains unclear how these senescent hepatocytes can trigger DR-driven regeneration and the potential role of the other non-parenchymal cells, such as the hepatic stellate cells or Kupffer cells.

We demonstrated that the LSECs phenotype plays a crucial role in degrading the laminin around DRs. When we prevented the capillarization of LSECs we did manage to slightly decrease the laminin content in the whole liver. However, such a decrease was not statistically significant (Fig. 4D). This was likely because LSECs are not the only cell types that can secrete or modify the ECM. The treated animals still have an ongoing CLD and other cell types, such as HSCs, KCs and BECs themselves can secrete the Laminin.

Additionally, we observed a decrease in the number of DRs. We suggested that having fewer capillarized LSECs creates a more favorable environment that pushes more DRs to differentiate into hepatocytes.

Our data showed that only preventing capillarization of the LSECs had an impact on DR-driven regeneration, suggesting that during CLD there is a switch from pro-regenerative to pro-fibrotic phenotype of the LSECs. It remains unclear whether capillarized CD34+ LSECs arise from a phenotypic switch of fenestrated CD34-LSECs or from the proliferation of vascular CD34+ endothelial cells. Further experiments are required to confirm such a hypothesis.

Taken together, our study uncovered the conditions associated with regenerative DRs and elucidated the underlying mechanism of regulation involving the CD34^+^ LSECs-LAM-DR microenvironment. Targeting this axis may assist in finding therapeutic strategies aimed at mitigating CLD and improving patient outcomes.

## Acknowledgments

D. Brusa from IREC FACS platform and Dr. Caroline Bouzin and Aurelie Daumerie from IREC 2IP platform for excellent technical assistance; Dr. Yves Horsmans and Dr. Shani Ben-Moshe for discussion and critical comments on the manuscript.

Fonds de la Recherche Scientifique - FNRS (1.B306.22, Rita Manco)

Fonds de la Recherche Scientifique - FNRS (FC36255, Maxime De Rudder)

Fonds de la Recherche Scientifique - FNRS (Alexandra Dili)

Foundation Saint Luc (PV)

PNRR MIUR DM 1061/2021 (DOT132670H-3, Camilla Moliterni)

## Author contributions

Conceptualization: RM, IAL; Methodology: RM, CM, GN, MDR, CP, LD, MF; Investigation: RM, CM, GN, MDR, AD, IAL; Visualization: RM, CM, IAL; Funding acquisition: RM, IAL Supervision: RM, IAL; Writing – original draft: RM, IAL.

## Data and materials availability

Single cell datasets used in this paper are already published. Mouse LSECs scRNAseq (Chromium, 10x Genomics) dataset were acquired from GSE147581 (n=3). The human MASH without cirrhosis was from ref. 28 with GSE202379. The human PSC dataset was from ref. 29 with GSE247128.

## Materials and Methods

### Animal models

To trace BECs/DR cells we used Osteopontin-iCreERT2 (OPN-Cre) mice crossed with Rosa26R^YFP^. To achieve Cre-LoxP recombination, tamoxifen (T5648; Sigma) at a concentration of 30 mg/ml corn oil was injected intraperitoneally at 100 mg/kg of bodyweight for 2 consecutive days on 21 and 23 days old OPN-Cre; Rosa26R^YFP^. For liver endothelial cell fate mapping, we used Cdh5-PAC-Cre^ERT2^ mice crossed with the Rosa^26R^ mTomato/mGFP reporter mice (called from now on Cdh5-CreXROSA^mT/mG^). In this strain, *Cre-LoxP* recombination was achieved by tamoxifen injection (100 mg/kg of body weight) i.p. for 3 consecutive days. In all mice, one month after tamoxifen treatment (to ensure complete tamoxifen wash-out) chronic liver injury was induced by repeated intraperitoneal injection of carbon tetrachloride (CCl_4_) 3 times per 6 weeks. The starting dose of CCl_4_ was 500 ml/kg, with a dose increase up to 800 ml/kg when animals gained weight. Livers were analyzed 72 hours after the last CCl_4_ injections. BAY 60-2770 was administrated orally daily with a dose of 1 mg/kg.

Mice were housed at 4–5/cage, maintained at a constant temperature of 22 °C, exposed at all times to a 12 h light/12 h dark cycle and had access to food and water ad libitum. Animal care was provided in accordance with the guidelines for humane care for laboratory animals in accordance with European regulations and in conformity with ARRIVE guidelines. The study protocol was approved by the university ethics committee for the use of experimental animals (A14GAEN2022).

### Human samples

Ten cases from an already collected series of biopsies from patients with clinical well-characterized and histopathological staged MASH were retrieved from the archives of the Institute of Pathology (CHUV, Lausanne, Switzerland).

### Cell culture and double-spheroids

Immortalized Normal Mouse Cholangiocyte (NMC; kind gift from T. Shimosegawa) cells were cultured in advanced DMEM/F12 reduced medium (gibco #12634-010) supplemented with 1,5% FBS (gibco #A3160801), 4mM L-glutamine and 1% penicillin/streptomycin, at 37°C in a 5% CO_2_ atmosphere. Twice a week, having reached 80% confluence, the cells were detached with trypsin and maintained at an exponential growth rate.

For the double spheroids, we used a modified protocol published by Lee, Russo et al(*18*). Briefly, columns containing mineral oil were formed using autoclaved 10µl pipette tips without filters. Collagen Type I, Rat Tail (Corning #354236) with 1.5% NaOH 1N was diluted to a final concentration of 1mg/ml using a solution composed of DMEM and filtered 10x reconstitution buffer containing HEPES and Sodium bicarbonate. Laminin (Corning #354232) was thawed on ice and used at a 3:1 ratio [three parts of Laminin and one part of Collagen I] once completely dissolved. Meanwhile, cells were detached, counted and resuspend to achieve a concentration of 50,000 cells per µl in either collagen or the laminin: collagen I, 3:1 v:v solution. 1µl of cells in the different extracellular matrix proteins was pipetted in the oil of the previously generated columns, and the mixture was incubated at 37°C for 1h30. Subsequently, cells were washed in FBS-free medium to remove oil residues, and cores spheroid were re-suspended in a second layer of matrix without cells and plated in a 96-well plate. After 1 h of incubation at 37°C, medium was added, and images were captured every day using a ZEISS LSM800 confocal microscope equipped with Zen Blue edition software, obtaining 20 images at 10µm intervals over 6 time points.

### Immunohistochemistry and immunofluorescence

Parts of each liver lobe were fixed in 4% formalin. Four µm formalin-fixed, paraffin-embedded liver sections were prepared. For immunohistochemistry detection, slides were deparaffinized and rehydrated; endogenous peroxidases immersed in 3% H2O2 in methanol for 15 minutes to be blocked, then the slides were submitted to heat-induced epitope retrieval in citrate buffer (pH 5.8-6.0) for 30 minutes at 100°C. Nonspecific binding sites were blocked incubating the slides for 1 hour in a solution of 10% BSA, 3% Milk and 0.3% Triton in PBS, following by incubation overnight at 4°C with primary antibodies (Details in Table S4). Detection was performed using either secondary Antibodies with AlexaFluor 488/ 594 or 647 fluorescent conjugates (nuclei were detected by incubation with DAPI) or with HRP conjugated secondary Antibodies, with subsequent DAB or TSA exposure.

For each case, two sequential new recuts of the pre-existing blocks were done to perform the following immunohistochemical analyses: 1. CK7 staining to identify the DR cells and newly formed hepatocytes, combined with Laminin, and 2. CK7 staining, combined with CD34. The double immunostainings were performed on a Ventana Discovery XT (Roche Diagnostics Switzerland) according to the manufacturer recommendations, starting with Laminin detection followed by CK7 detection, and by CD34 detection followed by CK7 detection, respectively. Antigen retrieval with Citrate based buffer was used, during 32 minutes for Laminin and CD34, and 8 minutes for CK7. All antibodies were incubated at 37°C for 32 minutes and revelation was perfomed with OmniMap anti-Rb HRP (760-4311) for Laminin and OmniMap anti-Ms HRP (760-4310) for CD34 and CK7. Laminin and CD34 are shown in red (Discovery Purple HRP kit) and CK7 is shown in green (Discovery Green HRP kit).

### RNA extraction and RT-qPCR

At sacrifice, parts of the liver were rapidly excised and snap-frozen in liquid nitrogen for gene expression analysis. Total RNA was extracted from 80-100 mg of tissue using TRIpure Isolation reagent (Roche Diagnostics, Vilvoorde, Belgium). Quantitative real-time polymerase chain reaction (qRT-PCR) was performed by AB StepOne Plus (Applied Biosystems Foster City, CA). All qPCRs were quantified using relative standard curves and normalized on the mean of RPL19 housekeeping gene. Results are expressed as fold expression relative to expression in the control group using ΔΔCt method. All samples were run in duplicate. The primer-pairs used are detailed in Table S2.

### RNA Extraction Spheroids

The medium was aspirated from the plate, and spheroids were washed three times with sterile PBS. Subsequently, Cell Recovery Solution was added, and the plate was incubated for 15 minutes on ice. The spheroids were then transferred to 0.5ml autoclaved and centrifuged at 1000 g for 1 minute. The remaining steps, including homogenization and extraction, followed the protocol of the RNeasy Plus Micro Kit (Qiagen #74034). RNA obtained was quantified and converted into cDNA using the Promega kit, which includes MasterMix 5x (#M531A), RNAsin Ribonuclease inhibitor (#N211B), and M-MLV RT RNase (H-) point mutant (#M368C).

### Isolation of liver cells and Flow cytometry analysis

Mice were anesthetized with 10 mg/kg Ketamine (Zoetis Manufacturing & Research) and 10 mg/kg Xylazine (EuroVet) dissolved in 1× PBS and injected i.p.. Livers from Cdh5-Cre; Rosa26R^mT/mG^ mice were washed to remove the blood perfusing through the vena cava 15 ml of pre-warmed to 37 °C EGTA buffer. Then, liver were dissociated using 10-15 ml of pre-warmed to 37 °C enzyme buffer solution (EBS) containing 0.4 mg/ml Pronase protease (Roche) and then with 15-20 ml of 37 °C EBS containing collagenase D (0.2 mg/ml) (Roche). Livers were extracted to a Petri dish with 25 ml of pre-warmed EBS and gently minced using forceps. Dissociated liver cells were filtered through a 100μm cell strainer. Two centrifugations of 30 g at 4°C for 3 min were performed to separate the non-parenchymal fraction from the hepatocytes. Supernatant was then centrifuged at 500g for 10min in 4°C. Pellet containing non parenchymal cells was resuspended in 1ml Red Blood Cell Lysis Buffer (Sigma), incubated at RT for 1min. EBS was then added and samples were centrifuged again in 500g for 10min in 4°C. Pellet was resuspended in 1-2ml of FACS buffer. Samples were then incubated with CD45, CD31, CD34, CXCR4, and CXCR7, according to the analyses.

### Molecular Cartography^TM^

Part of a lobule liver (caudate lobe) pieces were embedded in Tissue-Tek O.C.T.^TM^ Compound (Sakura) and snap frozen in isopentane (Sigma) chilled by liquid nitrogen. Embedded tissue pieces where stored at -80°C until cryosectioning. 10 µm liver slides were placed within capture areas on Resolve BioSciences slides. Samples were then sent to Resolve BioSciences on dry ice for analysis. Fixed samples were used for Molecular Cartography^TM^ (100-plex combinatorial single-molecule fluorescence in-situ hybridization) according to the manufacturer’s instructions (protocol 3.0; available for download from Resolve’s website to registered users). Probes used as detailed in Data S3.

Final image analysis was performed in Recognize software from Resolve BioSciences to examine specific Molecular CartographyTM signals.

### Single-cell RNA sequencing analysis

Mouse LSECs scRNAseq (Chromium, 10x Genomics) dataset were acquired from GSE147581 (n=3). The human MASH without cirrhosis was from the GSE202379. The human PSC dataset was from GSE247128. Analysis was performed with Seurat 5.0.1 in R 4.3.0 and R studio 2023.03.1.

### Mouse data

Variable genes were identified (FindVariableGenes, selection.method = “vst”, nfeatures = 2000). Cell clustering was based on PCA dimensionality reduction using the first 20 PCs and a resolution value of 0.2. We used cell type-specific markers to interpret the single cell clusters: Cdh5 for endothelial cells, Alb for hepatocytes, Cd3e for T-cells, Spp1 for cholangiocytes, Clec4g for macrophages, Dcn for hepatic stellate cells. We retained only cluster expressing Cdh5 and eGFP. We split the endothelial cells based on CD34 expression (threshold 1.5).

### Human data

We used the clustering of the original paper and select only cluster named with ‘LSEC’. Cell clustering was based on PCA dimensionality reduction using the first 5 PCs and a resolution value of 0.4. We split the endothelial cells based on CD34 expression (threshold 1.5).

**Table S1.**

Table with patients information.

**Table S2.**

Table of antibodies used with relative concentration.

**Table S3.**

Table of the RT-qPCR primers used.

