## Supplementary Figures for "Liver Sinusoidal Endothelial Cells and Laminin dictate cholangiocytes’ fate in chronic liver disease"

Supplementary Figure 1

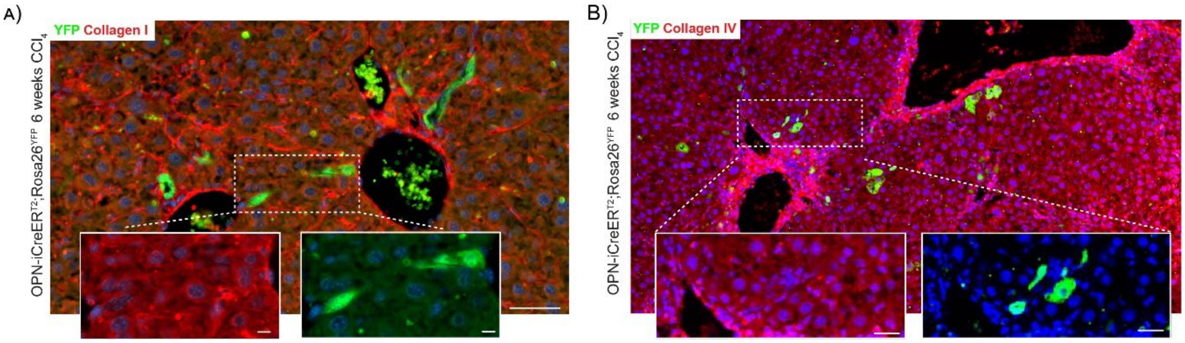


**Fig. S1. Collagen I and Collagen IV were not detected around DRs**

(A-B) Representative immunofluorescent images of Collagen I and Collagen IV (both in red) with YFP (Green) in liver section from OPN-iCreER$^{T2}$;Rosa26R$^{YFP}$ mice treated for 6 weeks with $CCl_4$.

### Supplementary Figure 2

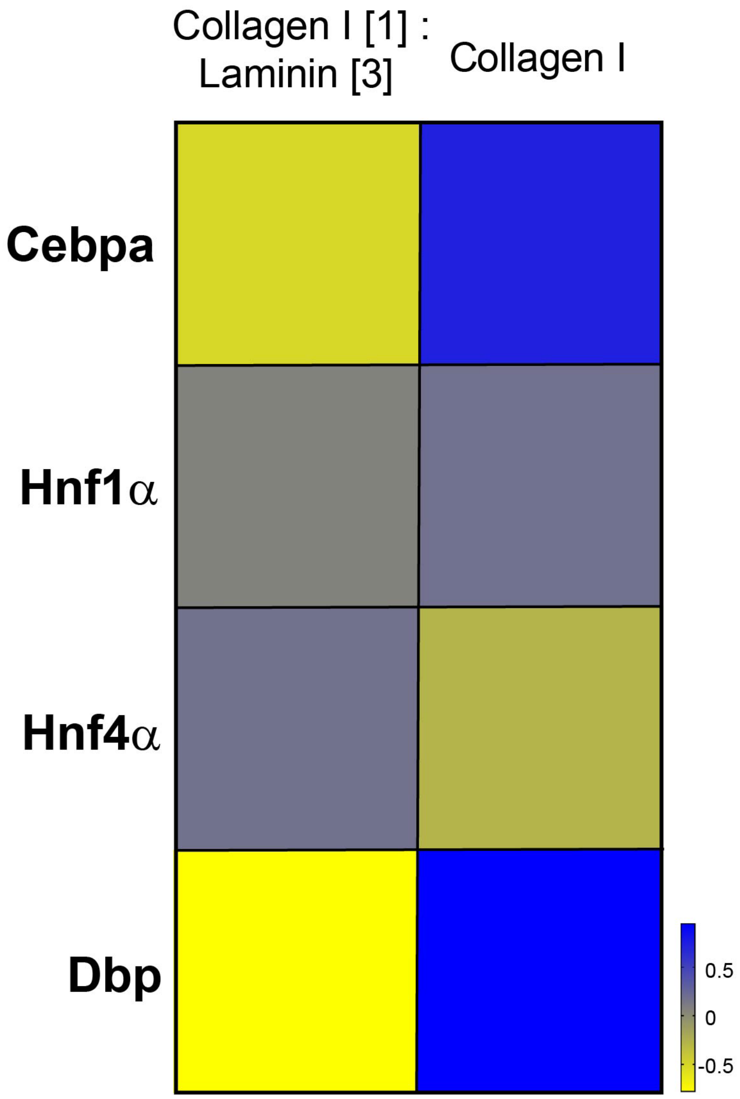


**Fig. S2. Cholangiocytes in contact with laminin have a lower expression of hepatocytes specific genes**

Heatmap of hepatocytes specific genes in cholangiocytes cultures in double-spheroids with or without laminin. Values are reported as z-score.

Supplementary Figure 3

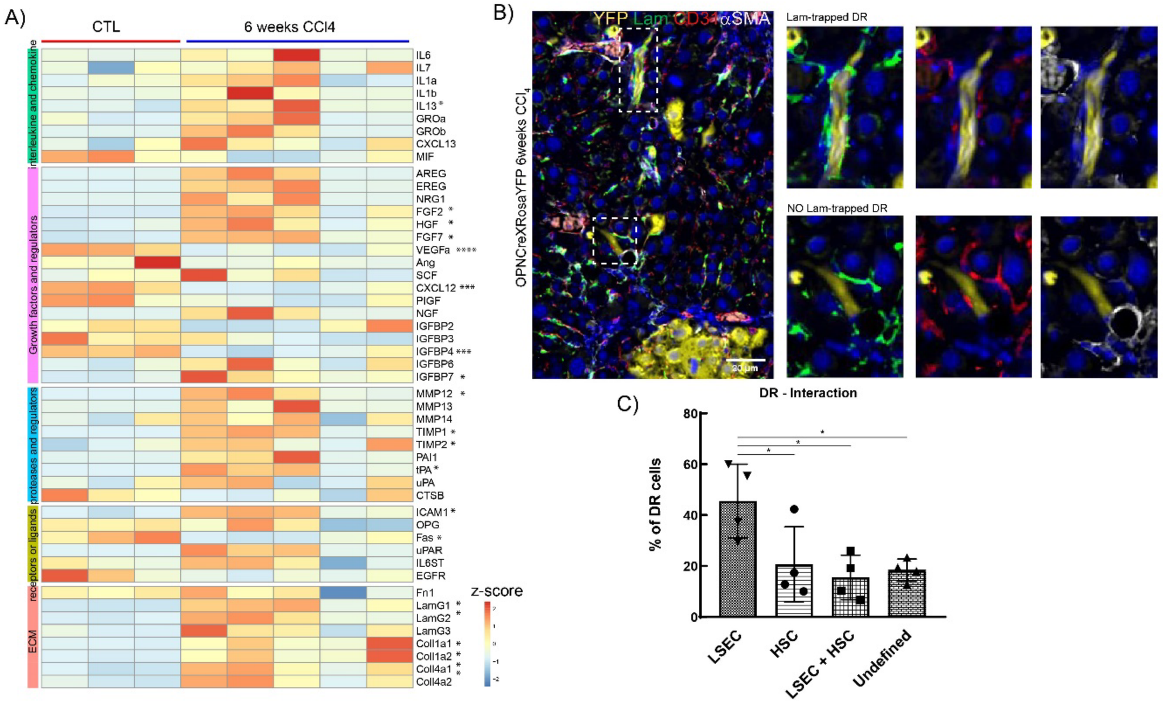


#### Fig. S3. During CLD, there is a dysregulation of SASPS angiogenic genes and LSECs are the cells closer to DRs.

(A) RT-qPCR of SASP represented by Z-score. (B) Representative multiplex immunofluorescent picture of liver section from OPNCreXRosaYFP animals treated with CCl4 for 6 weeks stained with YFP (yellow), Laminin (green), CD31 (red), and αSMA (gray). (C) quantification of the cell types near the DR cells. *P*-value was calculated by 1-Way ANOVA and Bonferroni post-test.

Supplementary Figure 4

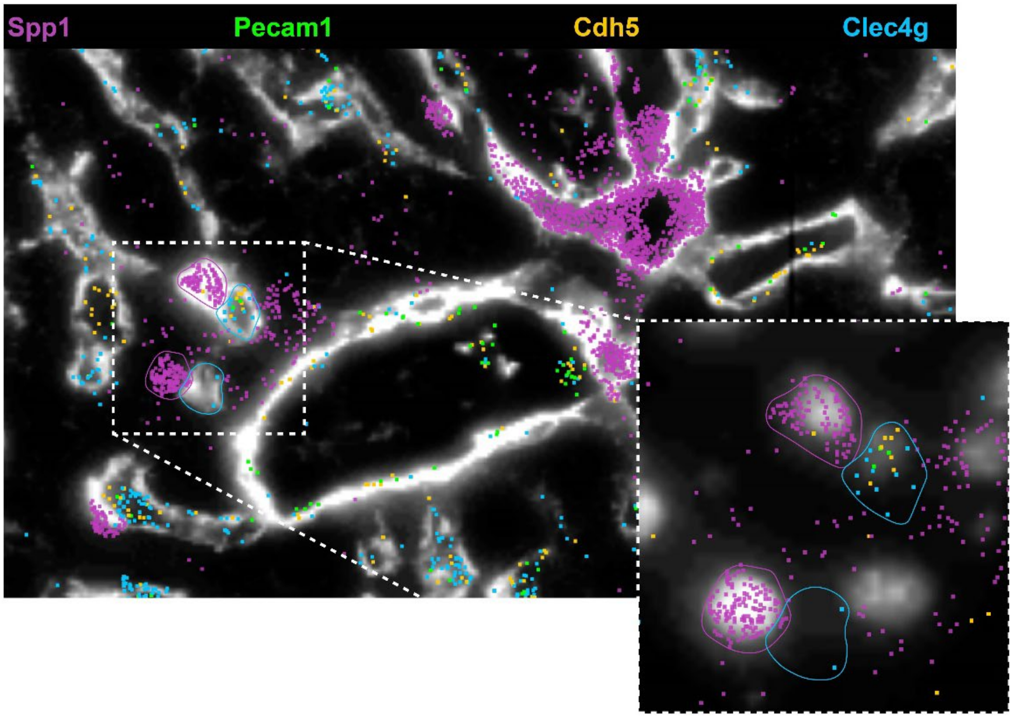


**Fig. S4. Molecular Cartography of indicated genes and IHC of laminin**

Supplementary Figure 5

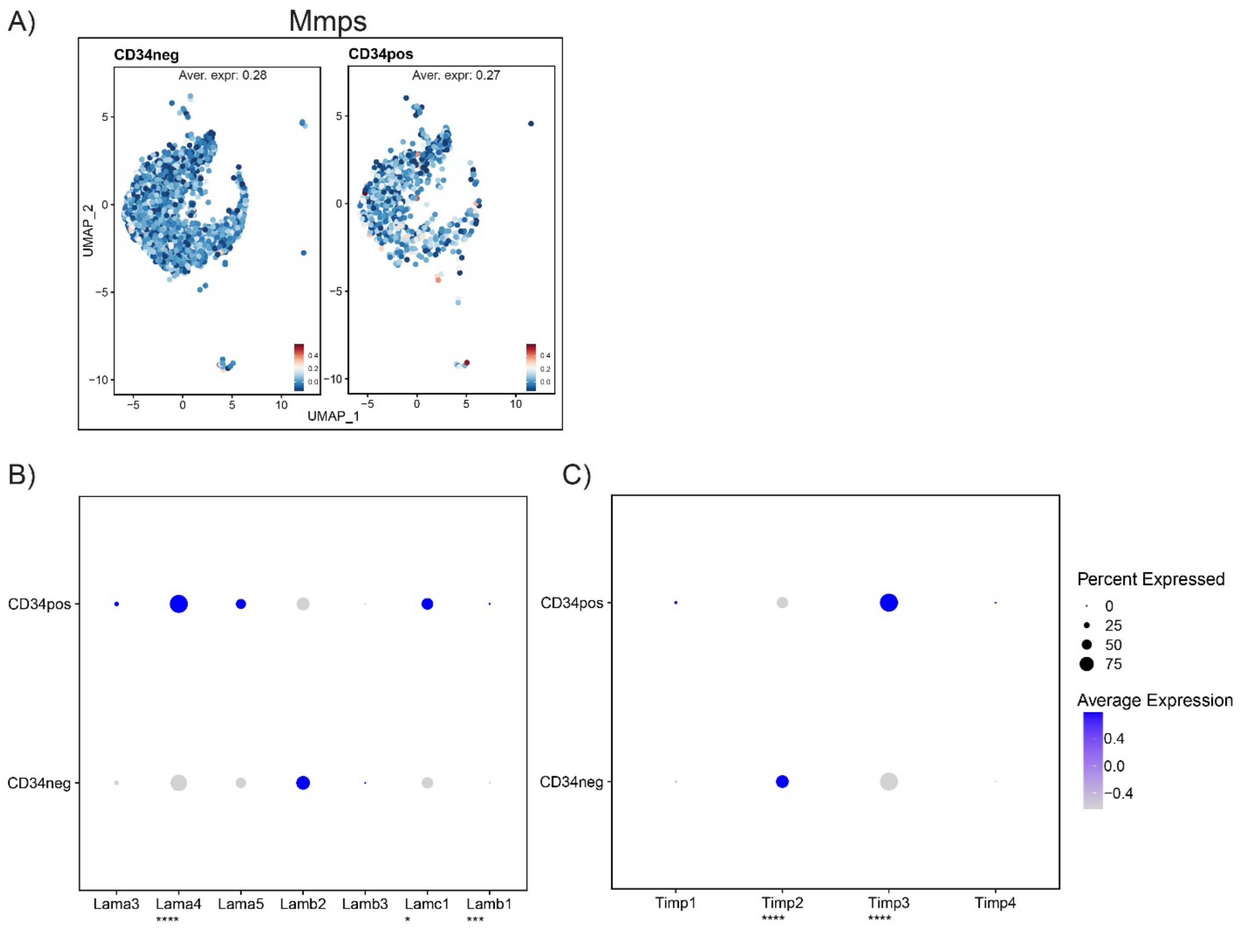


**Fig. S5. In CLD, CD34+ LSECs have the same expression of Mmps genes as CD34-, but increase the expression of Lama and Timp genes.**

(A) UMAP plots of LSECs sorted from CCl4-injured mice. (B-C) Dotplot of Lama and Timp genes in CD34+ and CD34- LSECs from CLD mice. Data are from Su et al., cmgh, 2021

### Supplementary Figure 6

A) B)

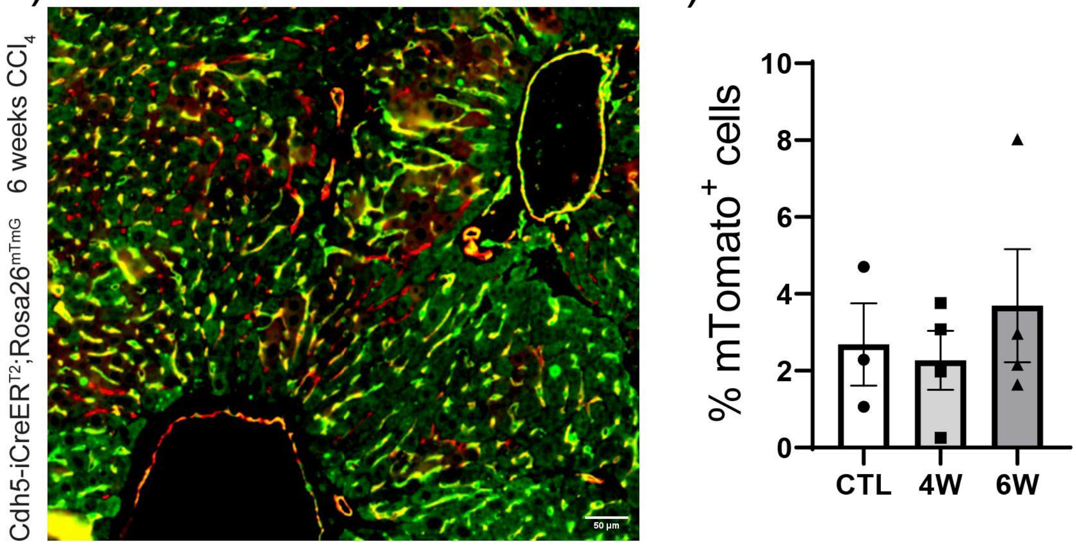


**Fig. S6. During CLD, there is no recruitment of endogenous LSECs.**

(A) Representative immunofluorescent image from liver of Cdh5-iCreER$^{T2}$;Rosa26$^{mTmG}$ mouse treated for 6 weeks with $CCl_4$. (B) FACS quantification of mTomato+ cells in Cdh5-iCreER$^{T2}$;Rosa26$^{mTmG}$ mice after 0 (CTL), 4weeks and 6weeks of $CCl_4$.

### Supplementary Figure 7

A)

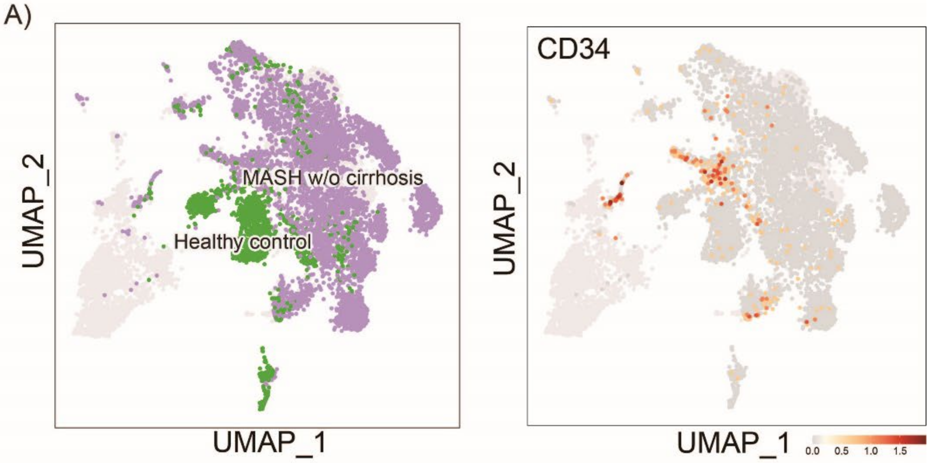


B)

| Group | % CD34⁺ endo cells |
|---|---|
| **Healthy Control** | 3.04 |
| **MASH w/o cirrhosis** | 4.50 |

C)

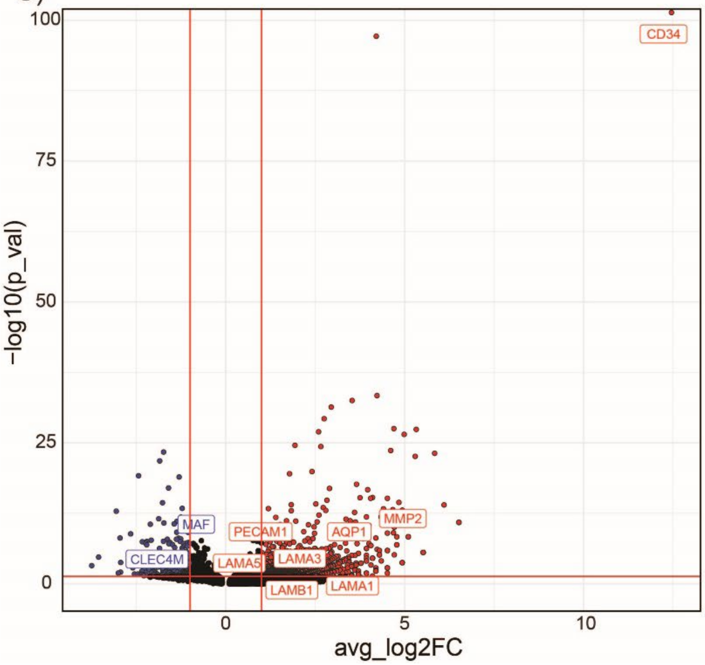


**Fig. S7. MASLD patients increase the expression of CD34+ and LAMA genes in LSECs.**
(A) UMAP plots showing the 2 groups we analyzed (Healthy control and MASH w/o cirrhosis) and the CD34+ expression. Plots were generated using the R Shiny app https://www.mohorianulab.org/shiny/vallier/LiverPlasticity_GribbenGalanakis2024. (B) Table with the % of CD34+ endo cells. (C) Volcano Plot showing the differentially expressed genes between CD34+ and CD34- LSECs.

Supplementary Figure 8

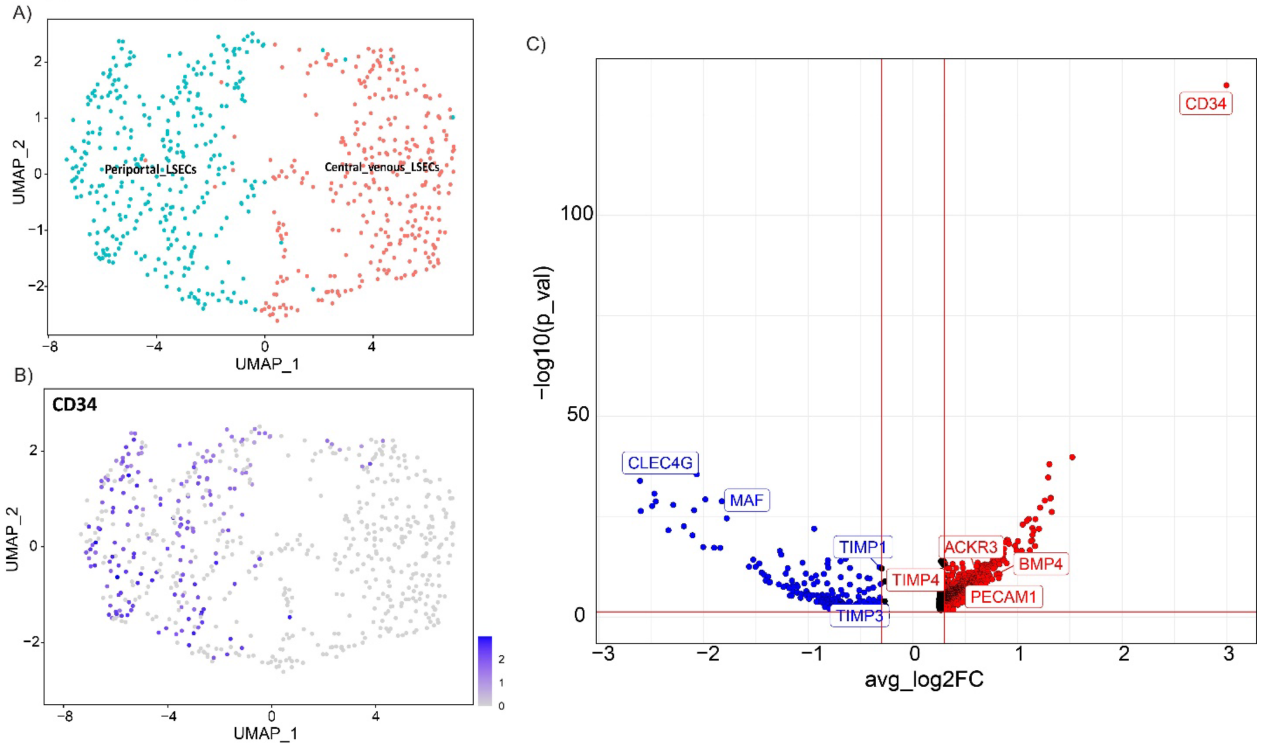


**Fig. S8. In PBCs patients, only periportal LSECs increase CD34 and LAM-remodeling specific genes expression.**

(A)UMAP showing the classification of the LSECs; (B) UMAP with CD34 expression. (C) Volcano Plot showing the differentially expressed genes between CD34+ and CD34- LSECs.
